# The importance of large-diameter trees in the wet tropical rainforests of Australia

**DOI:** 10.1101/474213

**Authors:** Matt Bradford, Helen T Murphy

## Abstract

Large trees are keystone structures in many terrestrial ecosystems. They contribute disproportionately to reproduction, recruitment and succession, and influence the structure, dynamics, and diversity of forests. Recently, researchers have become concerned about evidence showing rapid declines in large, old trees in a range of ecosystems across the globe. We used ≥10cm diameter at breast height (DBH) stem inventory data from 20, 0.5 ha forest plots spanning the wet tropical rainforest of Queensland, Australia to examine the contribution of large-diameter trees to above ground biomass (AGB), richness, dominance, mortality and recruitment. We show consistencies with tropical rainforest globally in that large-diameter trees (≥70 cm DBH) contribute much of the biomass (33%) from few trees (2.4% of stems ≥10cm DBH) with the density of the largest trees explaining much of the variation (62%) in AGB across plots. Measurement of AGB in the largest 5% of trees allows plot biomass to be predicted with ~85% precision. In contrast to rainforest in Africa and America, we show that a high proportion of species are capable of reaching a large-diameter in Australian wet tropical rainforest resulting in weak biomass hyperdominance (~10% of species account for 50% of the biomass) and high potential resilience to regional disturbances and global environmental change. We show that the high AGB in Australian tropical forests is driven primarily by the relatively high density of large trees coupled with contributions from significantly higher densities of medium size trees. Australian wet tropical rainforests are well positioned to maintain the current densities of large-diameter trees and high AGB into the future due to the species richness of large trees and a high density of replacement smaller trees.

## Introduction

Large trees are keystone structures in many terrestrial ecosystems including urban areas and agricultural systems [1]. They play a critical ecological role, storing large quantities of carbon, dominating canopies, providing food, shelter, habitat, and nesting cavities, and modulating microclimates and hydrological processes [2-5]. In forest ecosystems, large trees also contribute disproportionately to reproduction, recruitment and succession, and influence the structure, dynamics, and diversity of forests [6, 7]. Recently, researchers have become concerned about evidence showing rapid declines in large, old trees in a range of ecosystems across the globe [8]. Several reasons for this decline have been suggested including higher mortality rates in response to drought [9, 10] and cyclones [11], and the effects of fragmentation [12], logging, land clearing and agricultural intensification [2, 13].

Tropical forests make an important contribution to the global carbon cycle and the aboveground carbon balance of these forests is largely governed by the growth and mortality of individual trees [14]. Large trees in tropical forests have been shown to be particularly vulnerable to effects of fragmentation and lagged-mortality arising from damage sustained during logging activities [12, 15]. However, in tropical forests not subject to significant human-disturbance, evidence for decline in large trees is limited and long-term datasets are rare. Several authors have noted overall increases in mortality in trees across tropical forest plots in America and Asia though large trees have not been reported to be disproportionately affected [16]. Over 8.5 years, the mortality rate of trees >40m in height in lowland American rainforest was less than half the landscape-scale average for all canopy trees [5]. These authors suggest low mortality rates may be attributed to species-specific traits such as high wood density or delayed reproduction that increases survival through all life stages, increasing the probability of attaining tall heights. Alternatively (or in concert), low mortality in tall trees may be due to ecological advantages such as escape from physical damage from branch falls of lower stature trees or greater light interception which increases carbon gain [5]. It has been suggested [8] that elevated plant-growth rates in tropical forests, possibly resulting from rising atmospheric CO_2_, might result in larger numbers of large trees, particularly where other human disturbances are limited.

Large trees store large quantities of carbon and have been shown to drive variation in biomass in tropical forests across both the Neo- and Paleotropics [4, 17, 18]. Large trees (≥70 cm diameter at breast height (DBH) stored, on average, 25.1, 39.1 and 44.5% of above ground biomass (AGB) in South America, Southeast Asia and Africa, respectively, but represented only 1.5, 2.4 and 3.8% of trees >10 cm DBH (Table 1, [4]). Large trees also accumulate carbon much faster than smaller trees [19]. Large trees have been recently described as being ‘biomass hyperdominant’, that is, the functions of storing and producing carbon are concentrated in a small number of tree species [20]. For example, in tropical forest plot datasets from the Amazon and Africa, just ~1% [20] and 1.5% of tree species [17] were responsible for 50% of carbon storage and productivity. Understanding the dynamics of these large trees, and their response to changing environmental conditions, is clearly important for predicting the long-term functioning of tropical ecosystems as well as carbon storage and cycling.

**Table 1.** Large-diameter tree (>70 cm DBH) and AGB characteristics of Australian, Asian, American and African tropical rainforest.

|  | Mean AGB<br>(Mg/ha) | Number of large<br>stems/ha | Contribution of<br>large trees to<br>AGB (%) | Percentage of<br>large stems |
| --- | --- | --- | --- | --- |
| Australia<sup>#</sup> | 590.5±169.0 | 18.4±5.4 | 32.7 | 2.4 |
| Asia<sup>*</sup> | 393.3±109.3 | 13.4±6.7 | 39.1 | 2.4 |
| Amazon<sup>*</sup> | 287.8±105.0 | 7.5±5.3 | 25.1 | 1.5 |
| Africa<sup>*</sup> | 418.3±91.8 | 15.8±5.4 | 44.5 | 3.8 |

<sup>#</sup>This study
\*[4]

Australian wet tropical forests have among the highest biomass of tropical forests globally. On average, AGB in Australian lowland rainforest (<600 m) is 1.7 times higher than in lowland Amazonian forest, and between 1.2 and 1.3 times higher than in African and Asian lowland forests (Table 1, [4]). Interestingly, AGB values in Australian lowland forests are also considerably higher than those in Papua New Guinea [21], our nearest neighbour and most phylogenetically similar rainforest. Comparisons are similar or more pronounced in upland forests (600-1000 m) [22-24] and highland forests (1000-1500 m) [23, 25, 26]. We have monitored growth, recruitment and mortality of stems (≥10 cm DBH) in 20, 0.5 ha plots in the wet tropical rainforest of Australia for nearly 5 decades [27]. Here, we assess the contribution of large trees to carbon storage on our plots and examine changes in mortality and recruitment. We also discuss the role of large trees in the high biomass estimates seen in Australian wet tropical rainforest. We compare our results with those for tropical forests globally, highlighting convergent and divergent patterns. We demonstrate, 1) broad consistencies with other tropical rainforest globally confirming the importance of large-diameter trees to the carbon cycle, and 2) some divergence from other tropical rainforest globally, notably high species richness of large-diameter trees resulting in low biomass hyperdominance by species and some uncertainty in estimating AGB from the largest trees.

## Materials and Methods

### Study sites

The 20 CSIRO permanent study plots are situated in north-east Queensland, Australia, between 21.5°S, 149°E and 12.5°S, 143°E. The region is topographically diverse, and our dataset spans much of the geographical and environmental variation. The climate is tropical with mean annual rainfall ranging from 1200 mm to over 8000 mm on the higher coastal ranges. Seventeen of the plots are located within the Wet Tropics bioregion which consists of narrow coastal plains flanked by rugged mountains (to 1622 m) with extensive upland areas gradually sloping to the west. While covering only 0.24% of the Australian continent, the Wet Tropics region contains high levels of diversity and endemism of flora and fauna [28]. The plots were established between 1971 and 1980 in largely undisturbed forest and have been resurveyed every 2-15 years through to 2016. For a full description of the methodology and access to the data see [27]. At each census all stems ≥10 cm diameter are measured at DBH and mortality of stems ≥10 cm is recorded. Each individual is identified to species.

### Defining large-diameter trees

For moist forests in the Amazon with AGB of 85 - 400 Mg/ha, trees ≥70 cm diameter were identified as being important components of AGB [29]. Studies since have also defined large trees as ≥70 cm DBH [4, 18, 30], however others have used a definition that is specific to the particular study or forest type (e.g. >100 [31], >80 cm [32], 60-90 or >90 cm [15], and >60 cm [3]). In this study we define a large tree as being ≥70 cm DBH to allow for relevant pan-tropical comparisons.

### Above ground biomass estimates

Above ground biomass estimations are most accurate when they incorporate DBH, wood density and tree height [33] and we consider equation 2 described by Chave et al. [34] to be the most appropriate for AGB estimations in our forests. Unfortunately, height estimates for our plots were only collected at establishment and in 1998. Therefore, we assessed two methods of deriving height from DBH; 1) a pantropical equation that assumes a relationship between environmental stress and tree height [34], and 2) an Australian moist forest equation developed from height diameter relationships [31]. We compared these derived heights with our 1998 height data and heights collected from 22,694 trees at the Robson Creek 25 ha rainforest plot [35] also located within the Wet Tropics Bioregion. The first equation considerably underestimated measured height, and the second equation overestimated but approximated measured height (S1 and S2 Figs). We used the derived heights from both sources in equation 2 in [34] and compared the resulting AGB estimations to those using the actual measured heights from the 1998 and Robson Creek 25 ha data (S3 Fig). Estimations from both sources were less accurate than simply using an equation for tropical moist forests without height [33]. The latter is therefore used in this study:

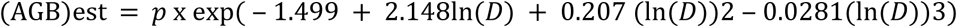

Where *p* = wood density (g/cm^3^) and *D* = DBH (cm). Wood density values were taken from a database compiled from the Australian literature and field collections. Values from the literature were used if sourced from northern Australia. Where more than one measurement was available, mean values were taken for a species. Where a species value was not available (n=19), the genus mean was used (n=14). Where a genus mean was not available, the family mean was used (n=4). Where a family mean was not available, the plot mean was used (n=1).

### Data analysis

Trees in each plot were ranked by decreasing size according to their AGB and their contribution to total AGB calculated. We calculated the number of species that collectively account for 50% of the total biomass both at the plot and regional scale at the most recent survey. The contribution of the largest trees to total species richness for each plot was calculated.

We used linear regression to assess the variation in total AGB explained by the single largest tree in each plot, and by the top 5, 10, 15 and 20% of largest trees in each plot. Relative root mean square errors were calculated to assess precision of the regression model.

Mortality and recruitment rates were calculated as per Condit, Ashton (36). Thus, for mortality:

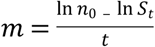

For recruitment:

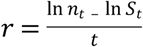

Where the census interval is *t, n*_o_ is the population size at time 0 and *n*_t_ is the population size at time *t*. The number of survivors at time *t* is *S*_t_. Mortality and recruitment rates were calculated for each size class (large trees ≥70 cm DBH, medium trees 30-70 cm, and small trees <30 cm DBH) at each census interval for each plot and then averaged by decade across all plots.

## Results

Over all census periods, 81 species were recorded as large trees (≥70 cm), which is 16.6% of all species in the dataset. Three species were strangler figs (*Ficus* sp.: Moraceae). The family Myrtaceae had the highest number of large tree species (n=11), while the Sterculiaceae had the highest proportion of species reaching large diameter status (42%, n=5). Species that grew into large trees had a significantly lower wood density (mean = 0.59 g/cm^3^) than species that did not (mean = 0.64 g/cm^3^) (ANOVA F_(1, 488)_ = 6.51, P = 0.011).

Across the 20 plots the mean AGB at the last census was 590 ± 169 SD Mg/ha (range 307 – 909) (Table 2). The size class 10 −20 cm DBH contributed 60.5% of the trees (Fig 1). At the last census the size class 30 - 40 cm DBH contributed the most AGB of any 10 cm size class bracket (13.7%), a shift from the first census where the 40 – 50 cm DBH size class contributed the most (15.0%) (Fig 2). The total number of large trees was 169 at the first survey and 182 at the last survey. At the last census the average number of large trees per hectare was 18.4 ±5.4 SD, comprising 2.4% of total trees, and large trees accounted for 32.7% of AGB across all plots (range 0 – 52.3%). The mean proportion of the total AGB accounted for by the cumulative number of largest trees increased rapidly reaching an average of 49% for the 20 largest trees (5% of the trees) and 84% for the largest 100 trees (27% of the trees) (Figs 3a and 3b).

**Table 2.** Above ground biomass and stem demographics of the 20 CSIRO permanent plots.

| Plot | First census |  |  |  |  | Last census |  |  |  |  |
| --- | --- | --- | --- | --- | --- | --- | --- | --- | --- | --- |
|  | Date | AGB | Number | Number | Largest | Date | AGB | Number | Number | Largest |
|  |  | (Mg/ha) | of stems | of stems | stem |  | (Mg/ha) | of stems | of stems | stem |
|  |  |  |  | ≥70 cm | (cm DBH) |  |  |  | ≥70 cm | (cm DBH) |
|  |  |  |  | DBH |  |  |  |  | DBH |  |
| ep18 | 1973 | 825 | 454 | 11 | 125.7 | 2016 | 838 | 427 | 13 | 120.5 |
| ep19 | 1975 | 480 | 400 | 7 | 90.9 | 2016 | 307 | 387 | 3 | 87.8 |
| ep2 | 1971 | 296 | 462 | 0 | 54.8 | 2015 | 343 | 511 | 0 | 55.4 |
| ep29 | 1975 | 429 | 494 | 1 | 83.7 | 2015 | 478 | 409 | 2 | 98.6 |
| ep3 | 1971 | 782 | 502 | 12 | 113.1 | 2015 | 814 | 466 | 15 | 118.0 |
| ep30 | 1976 | 680 | 552 | 8 | 94.9 | 2013 | 674 | 550 | 10 | 94.2 |
| ep31 | 1976 | 667 | 238 | 13 | 108.9 | 2013 | 537 | 191 | 6 | 118.7 |
| ep32 | 1975 | 394 | 447 | 4 | 106.6 | 2015 | 421 | 402 | 5 | 119.0 |
| ep33 | 1976 | 743 | 315 | 13 | 196.5 | 2015 | 806 | 250 | 23 | 122.7 |
| ep34 | 1976 | 584 | 302 | 10 | 118.1 | 2016 | 530 | 259 | 10 | 122.9 |
| ep35 | 1977 | 445 | 501 | 2 | 84.5 | 2016 | 551 | 399 | 5 | 92.2 |
| ep37 | 1977 | 978 | 386 | 14 | 147.7 | 2013 | 909 | 440 | 12 | 158.8 |
| ep38 | 1977 | 513 | 382 | 6 | 131.2 | 2016 | 505 | 324 | 7 | 134.4 |
| ep4 | 1972 | 369 | 486 | 1 | 70.4 | 2016 | 456 | 454 | 6 | 87.0 |
| ep40 | 1978 | 697 | 496 | 9 | 106.6 | 2013 | 627 | 414 | 9 | 120.6 |
| ep41 | 1977 | 519 | 395 | 3 | 88.4 | 2015 | 589 | 314 | 5 | 77.8 |
| ep42 | 1977 | 547 | 243 | 17 | 134.8 | 2013 | 403 | 234 | 8 | 126.5 |
| ep43 | 1978 | 730 | 387 | 16 | 127.3 | 2016 | 635 | 349 | 15 | 116.0 |
| ep44 | 1980 | 675 | 439 | 11 | 109.3 | 2013 | 708 | 399 | 12 | 111.5 |
| ep9 | 1972 | 638 | 441 | 11 | 121.2 | 2015 | 679 | 411 | 16 | 108.1 |
| Mean±SD |  | 599±172 | 416±87 | 8.5±5.2 |  |  | 590±169 | 379±93 | 9.1±5.6 |  |

**Fig 1.**
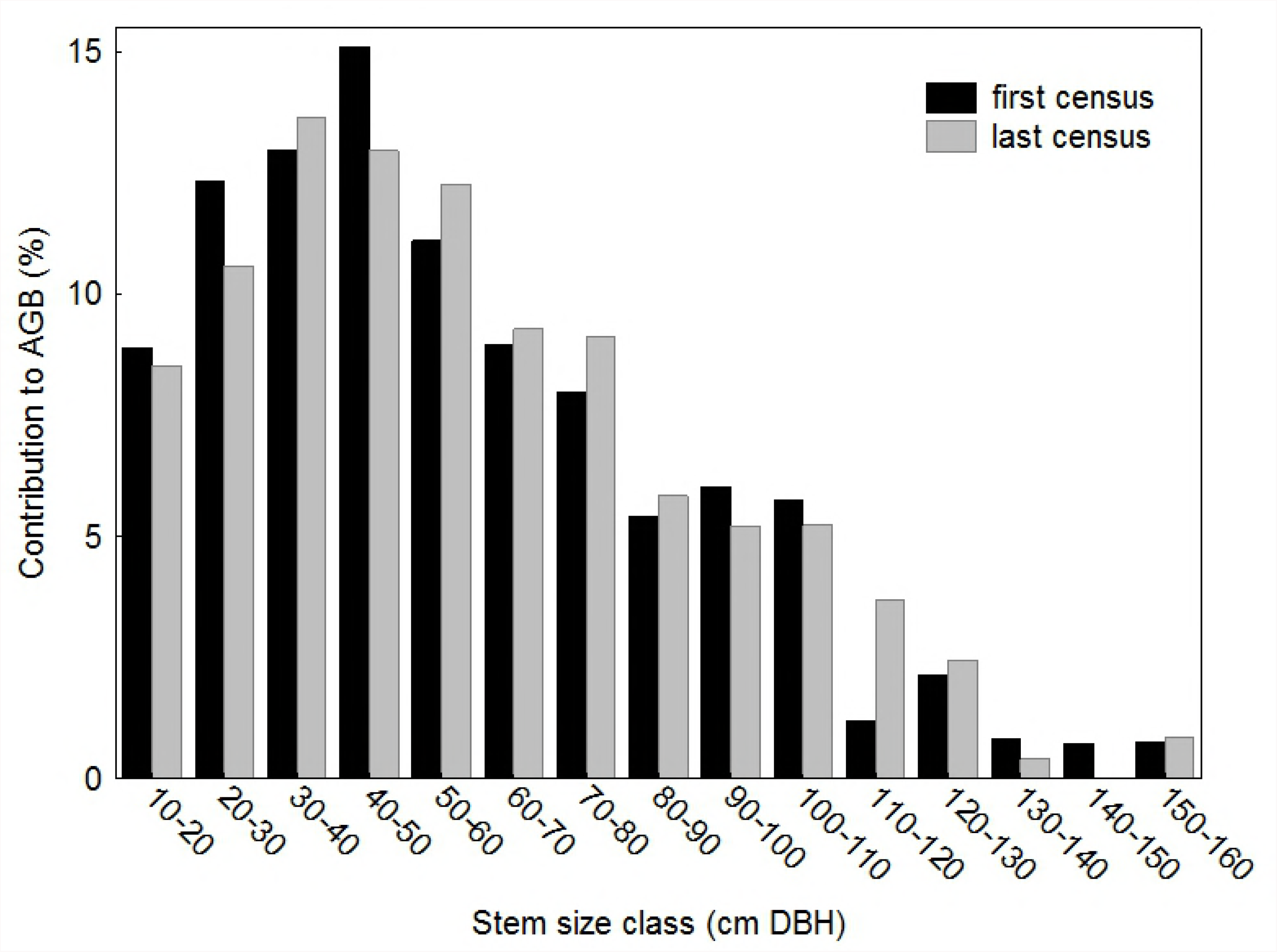
Contribution to total trees number by size class across the 20 CSIRO permanent plots. Note the broken Y-axis.

**Fig 2.**
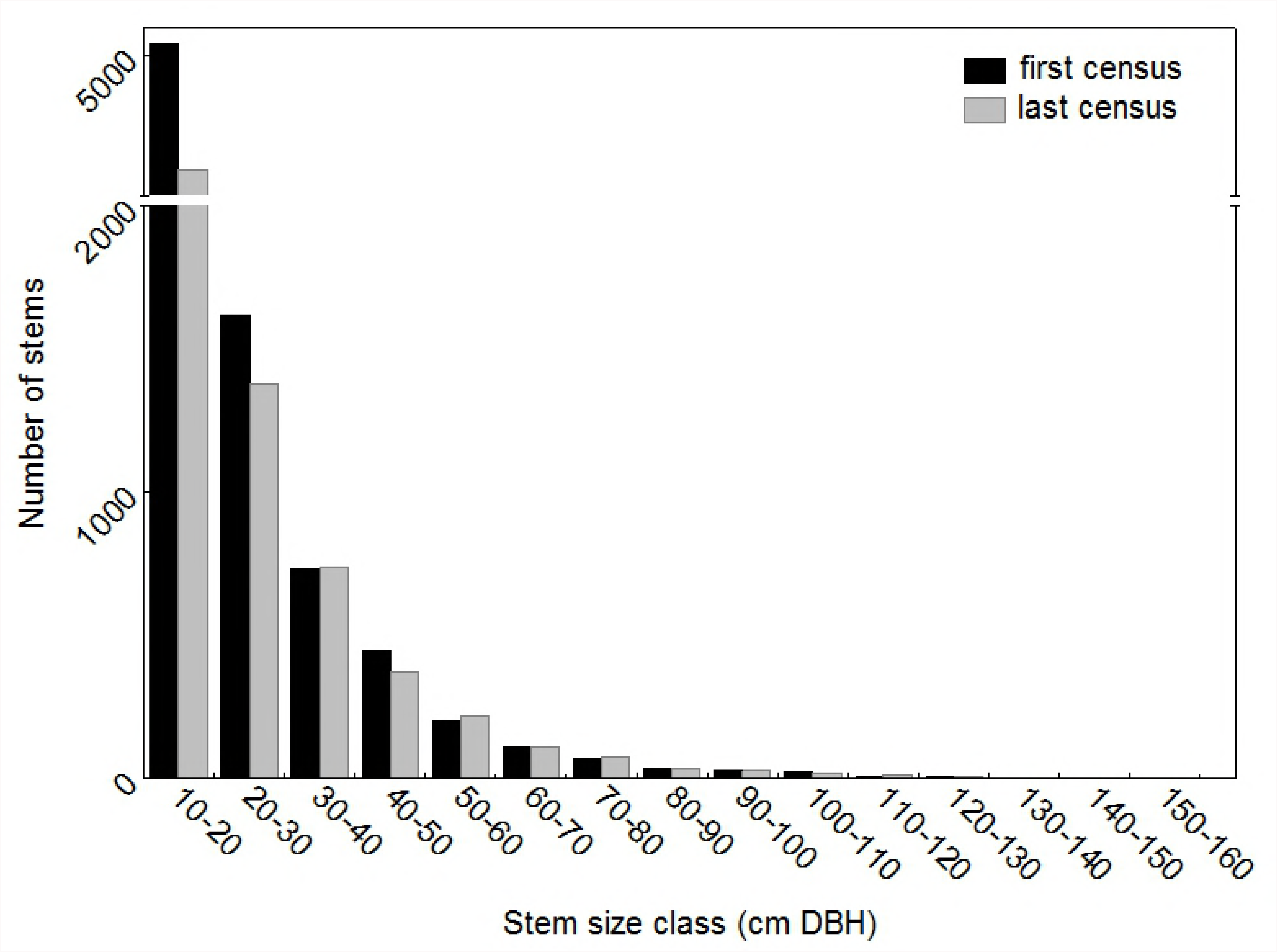
Contribution to total AGB by size class across the 20 CSIRO permanent plots.

**Fig 3.**
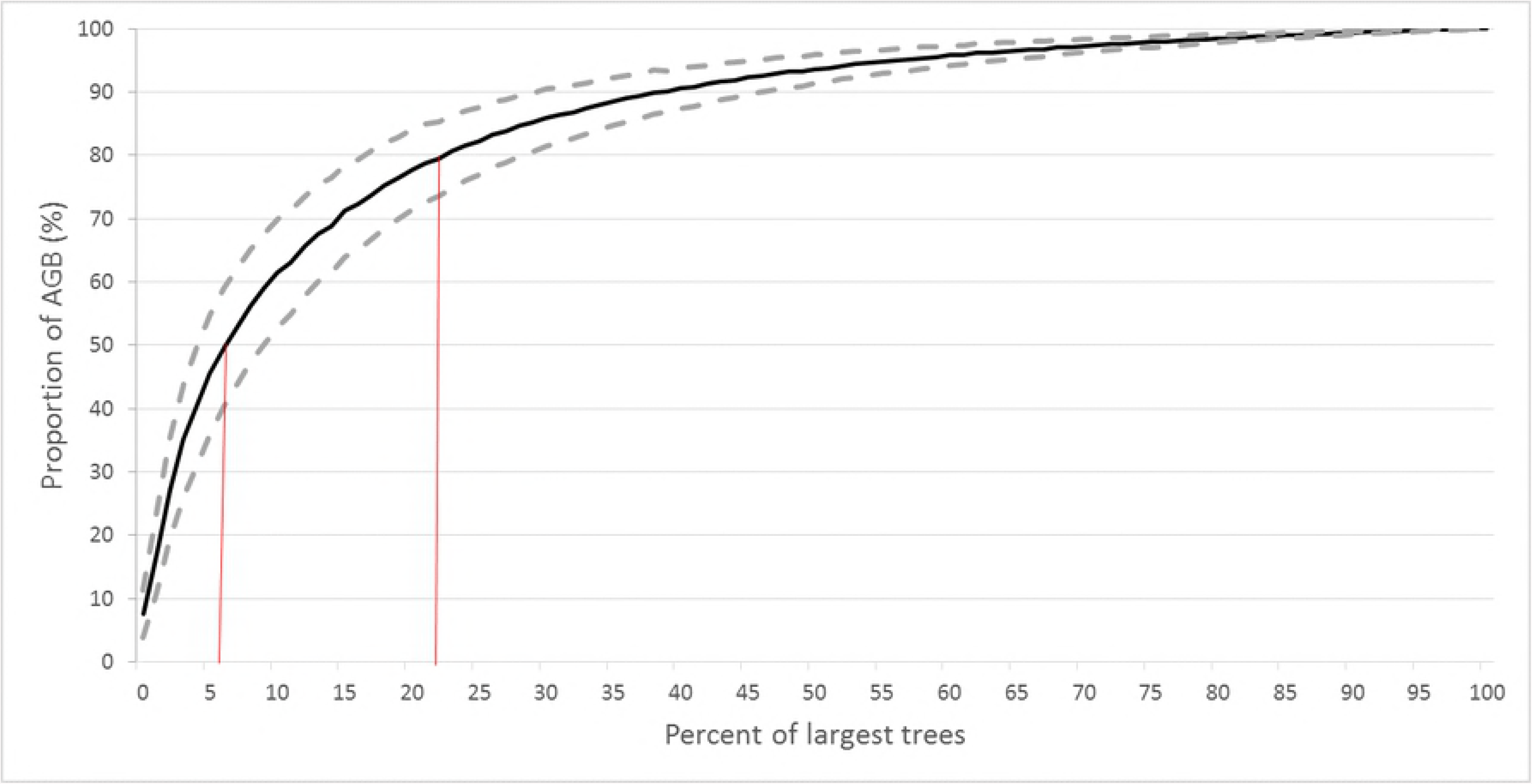
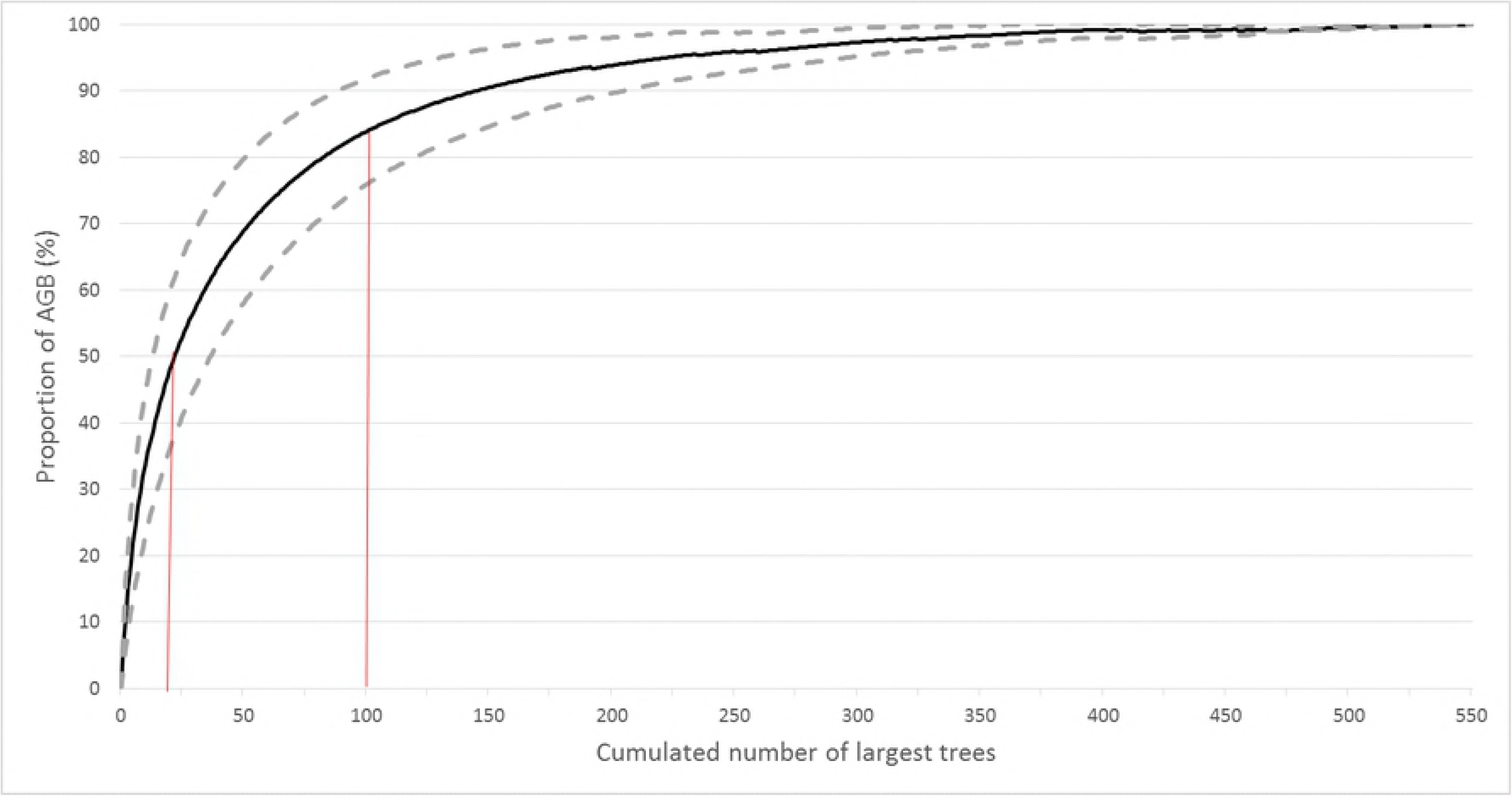
Proportion of AGB accounted for by the largest trees. (a) percent of largest trees and (b) cumulative number of largest trees. Vertical lines indicate the (a) percent of largest trees accounting for 50% and 80% of total AGB and (b) the proportion of AGB accounted for by the 20 and 100 largest trees. The dashed lines represent ± 1 SD of the mean.

The number of trees ≥70 cm DBH explained 62% of the variation in AGB across plots. The AGB of the single largest tree in each plot explained 25% of the variation in total AGB across all plots and the AGB of the top 5% of largest trees explained approximately 70% (Table 3).

**Table 3.**
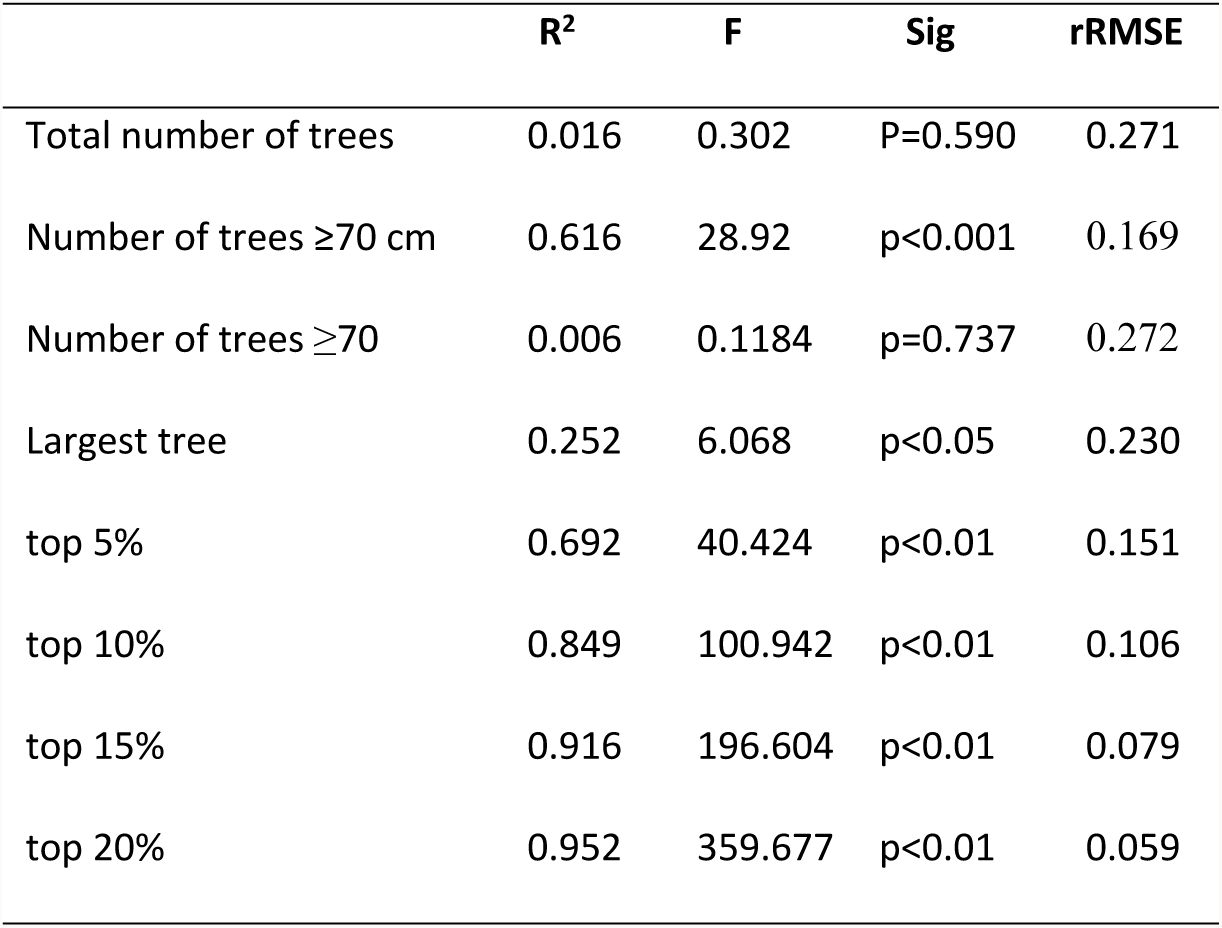
Linear regression of plot AGB against number of trees per plot, AGB of the largest tree per plot, and the largest 5, 10, 15 and 20% of trees in each plot.

Species richness was relatively high among the largest trees with the top 25 largest trees in a plot on average accounting for nearly 25% of total species richness in that plot; the 100 largest trees in a plot accounted for 56% of total species richness (Fig 4). At the last census, 123 species out of 443 (27.8%) contributed to the top 50% of total AGB across all plots. At the plot level, an average of 20% of species contributed to the top 50% of AGB in the last census (range 3% to 44%). The mean DBH for biomass hyperdominant species was 74.7 cm ± 25.5 SD compared with non-biomass hyperdominant species at 39 cm ± 27 SD.

**Fig 4.**
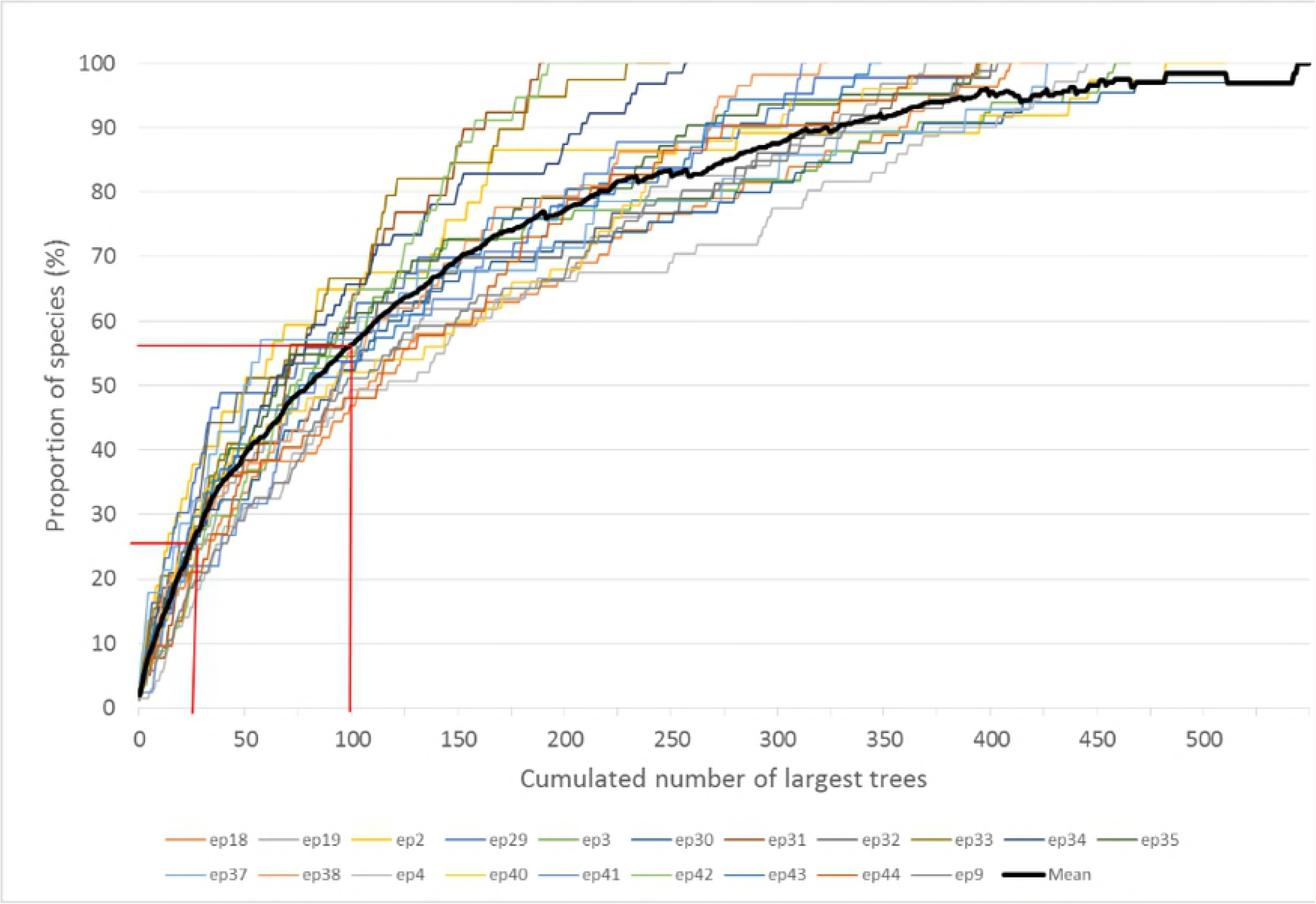
The proportion of total species accounted for by the cumulated number of largest trees. Results are displayed for each individual plot (coloured lines) with the heavy black line showing the mean. Vertical lines indicate the proportion of species accounted for by the top 25 and top 100 largest trees.

The mean annual rate of mortality for large trees was higher than for small and medium size trees in the first and last decade (decades beginning 1970 and 2010) and lowest in the decade beginning 1990 (Fig 5). Recruitment of large trees was also lowest in the 1970s but was higher than that for small and medium trees for the remainder of the monitoring period, though variation was high due to smaller overall numbers.

**Fig 5.**
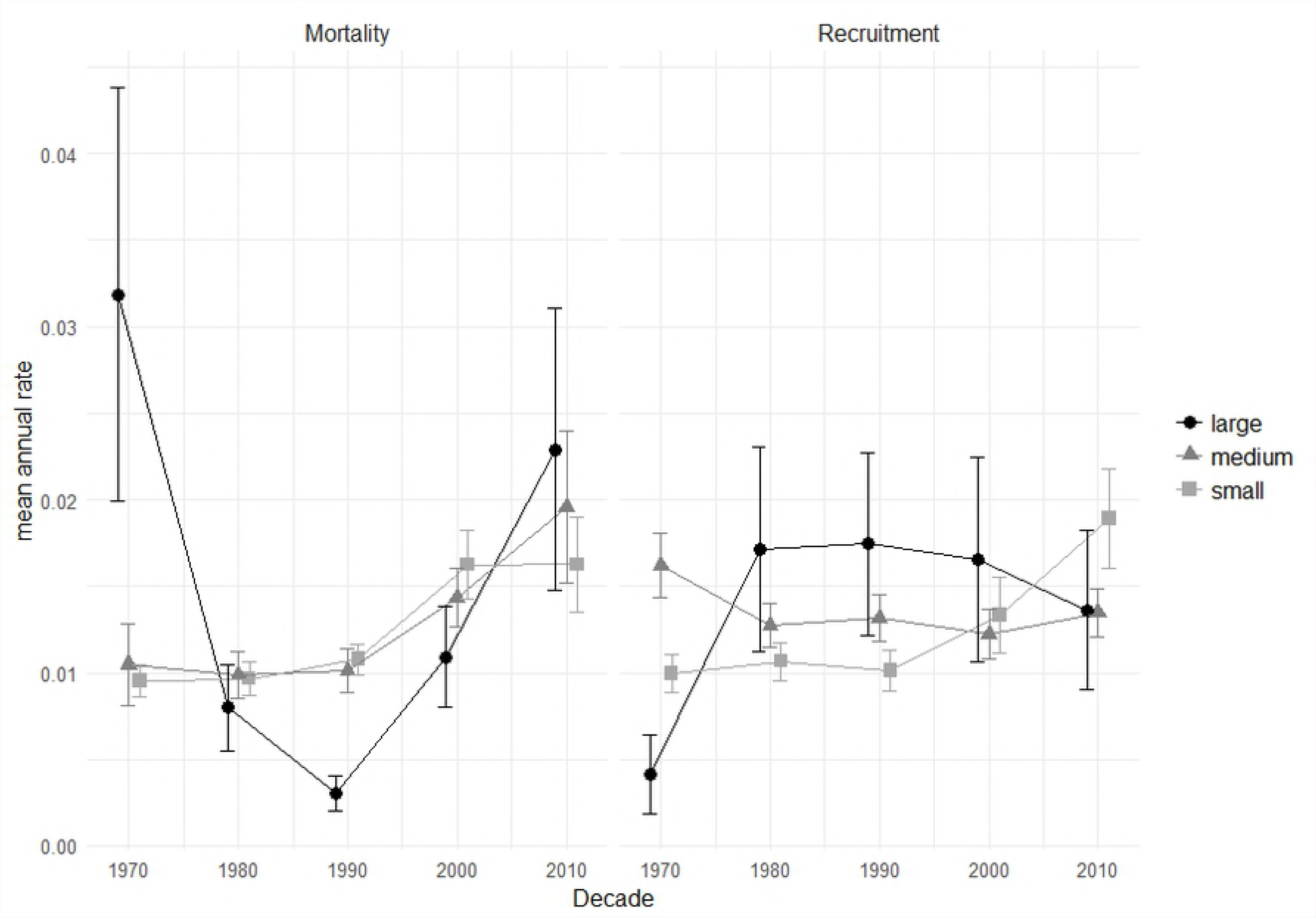
Mean annual rate of mortality and recruitment across all plots during each decade for large (≥70 cm dbh), medium (30-70 cm dbh) and small (<30cm dbh) trees.

## Discussion

### The contribution of large trees to above ground biomass

The role of a small number of large trees in driving forest biomass is now well recognised [3, 4, 18] and the concentration of AGB in a limited number of large trees has been quantified recently across the tropics [4, 17, 18, 37]. Despite Australian wet tropical rainforest holding considerably more biomass than rainforests elsewhere in Asia, Africa and America, there are consistencies in the contribution of the largest trees to biomass. Large-diameter trees (≥70 cm DBH) contribute approximately 33% of biomass and comprise 2.4% of trees ≥10 cm DBH in Australian wet tropical rainforest; within the range of values reported for Asian and African forests but significantly greater than reported for Amazonian forests (Table 1, [3]). The density of the largest trees explains much of the variation (~62%) in AGB across the plots, slightly less than the average pantropical estimate of ~70% (excluding Australia) [4].

However, the high biomass and stem density of Australian wet tropical rainforest results in some inconsistencies with pan tropical rainforests in total AGB prediction from the largest trees. The AGB of the single largest tree on each of our plots only explained 25% of the variation in AGB total across all plots which is approximately half that in African forests [17] and in North and South American forests [37]. The largest 5% of trees in a plot (ranging from 9 to 27 trees, average 18.5) explains 70% of the variance across plots, lower than that for African forests where the largest 20 trees explain 87% of variance [17]. In our plots measurement of the top 10 to 15% of trees (average 38 to 58 trees) is needed to explain close to 90% of the variation in AGB across plots. Measurement of AGB for the top 5% of trees allows an estimate of AGB total with approximately 85% precision (table 3), similar to that reported for African forests [17] and slightly better than reported in a global tropical forests analysis (excluding Australia) (~82% for the top 20 trees) [18].

We have shown [38] that the mean relative change in AGB in Australian wet tropical rainforest shifted from predominantly positive to predominantly negative during the 40 year monitoring period. Although the number of large trees across all plots in our current study increased by 8% over the census period we saw a recent increased mortality and decreased recruitment of large trees supporting a general trend of declining growth rates in Australian wet tropical rainforest. However, this must be viewed with caution as small overall numbers of large trees in 0.5 hectare plots (mean = 9.1± 5.6 SD) contribute considerable variation in rates of mortality and recruitment. In addition, productivity in Australian wet tropical rainforest is primarily influenced by large scale disturbance events [38]. Mortality and recruitment over the five decades of census was driven by three severe cyclones (1986, 2006 and 2010) and an extended dry period around 1986 that impacted all but three plots and may not reflect trends over the longer term.

### Biomass hyperdominance in large trees

Australian wet tropical rainforest does not appear to have strong biomass hyperdominance at the species level. Nearly 28% of species accounted for the top 50% of biomass across our plots compared with 1.5% for African forests plots [17] and 5.3% for Amazonian basin plots [20]. At the plot level, biomass hyperdominance ranged from 3% to 44% (average 20%); again much higher than the average 4.4% across African plots. A more realistic estimate for Amazonian-wide hyperdominance was suggested as ~1% considering the estimated 16,000 tree species that occur there [20]. We have far fewer plots in our dataset than those used in the Amazonian study, however the wet tropical rainforest of Australia is far less extensive and our dataset spans much of the geographical variation and environmental gradients across the region and includes ~31% of all species ≥10 cm DBH in the region. Australian wet tropical rainforest covers 10 000 km^2^ of the Australian continent (compared with 5.3 million km^2^ of Amazonian rainforest), and has an estimated 1450 tree species. If we consider our biomass hyperdominants are a reasonable representation of the region as a whole, then ~10% of tree species contribute 50% of the carbon stock in Australian wet tropical rainforest.

While examples of biomass hyperdominance are numerous in woodland communities in tropical Australia, examples in wet tropical rainforest are harder to find. In our dataset, *Backhousia bancroftii* accounts for the entirety of the top 50% of biomass in plot ep31, in part due to the species being less susceptible to cyclone damage than other species in the community [39]. Other less extensive examples of hyperdominance in the study area not represented in our plots are *Leptospermum wooroonoorum* that is restricted to wet exposed mountain ridges, *Ceratopetalum virchowii* that dominates on a particular low nutrient soil, and *Alstonia scholaris* that resists frequent cyclone disturbance allowing it to dominate on some exposed coastal slopes.

The relatively high diversity of species reaching a large size and contributing to biomass in Australian wet tropical rainforest has significant implications for the ongoing resilience of these forests. The loss of a single species or a group of closely related dominant species is unlikely to have the same consequences for forest carbon storage and forest function as it would in African or Amazonian forests. Due to the relatively small extent of our rainforest, a disturbance event such as a severe cyclone or a regional drought has the potential to impact a large proportion of the area. Having a high diversity of species in the largest size classes affords a greater level of resilience to such an event as Australian wet tropical rainforest species display a range of responses to disturbance and varied rates of recovery [39]. In addition, high taxonomic diversity safeguards against factors that target particular taxa such as the introduced fungal pathogen Myrtle rust that only infects the family Myrtaceae [40] and soil borne pathogens such as *Phytophthera* spp. that cause higher mortality in large trees and species in the family Elaeocarpaceae [41].

### Accounting for high AGB in Australian wet tropical rainforest

The high AGB in Australian wet tropical rainforest is largely a result of the high density of large-diameter trees with the density of trees ≥70 cm DBH explaining ~62% of the variation in AGB across plots. Australian wet tropical rainforest also have a significantly higher density of total (≥ 10 cm DBH), small, and medium trees than forests in Africa, South-east Asia and America (Fig 6). Although trees <70 cm DBH are not a good predictor of AGB in our plots (R^2^ = 0.0065, Table 3), medium sized trees (30-70 cm) contribute close to 50% of our total AGB and are seen as important contributors to AGB in some forests across the tropics [18]. The high density of small and medium sized trees in Australian wet tropical rainforest is likely due to the high frequency of large scale disturbance by tropical cyclones [38] which initially result in increased mortality but subsequently allow recruitment and growth into small and medium size classes. This is also presumably why we rarely see emergent trees in Australian wet tropical rainforests. There is also strong evidence that water use efficiency is much higher in Australian wet tropical rainforest than in similar forests globally resulting in trees rarely becoming water limited [42], most likely due to species evolving to survive in the generally dry continent of Australia. This potentially allows greater production of AGB presuming variables such as soil nutrients and solar radiation interception are not limiting.

**Fig 6.**
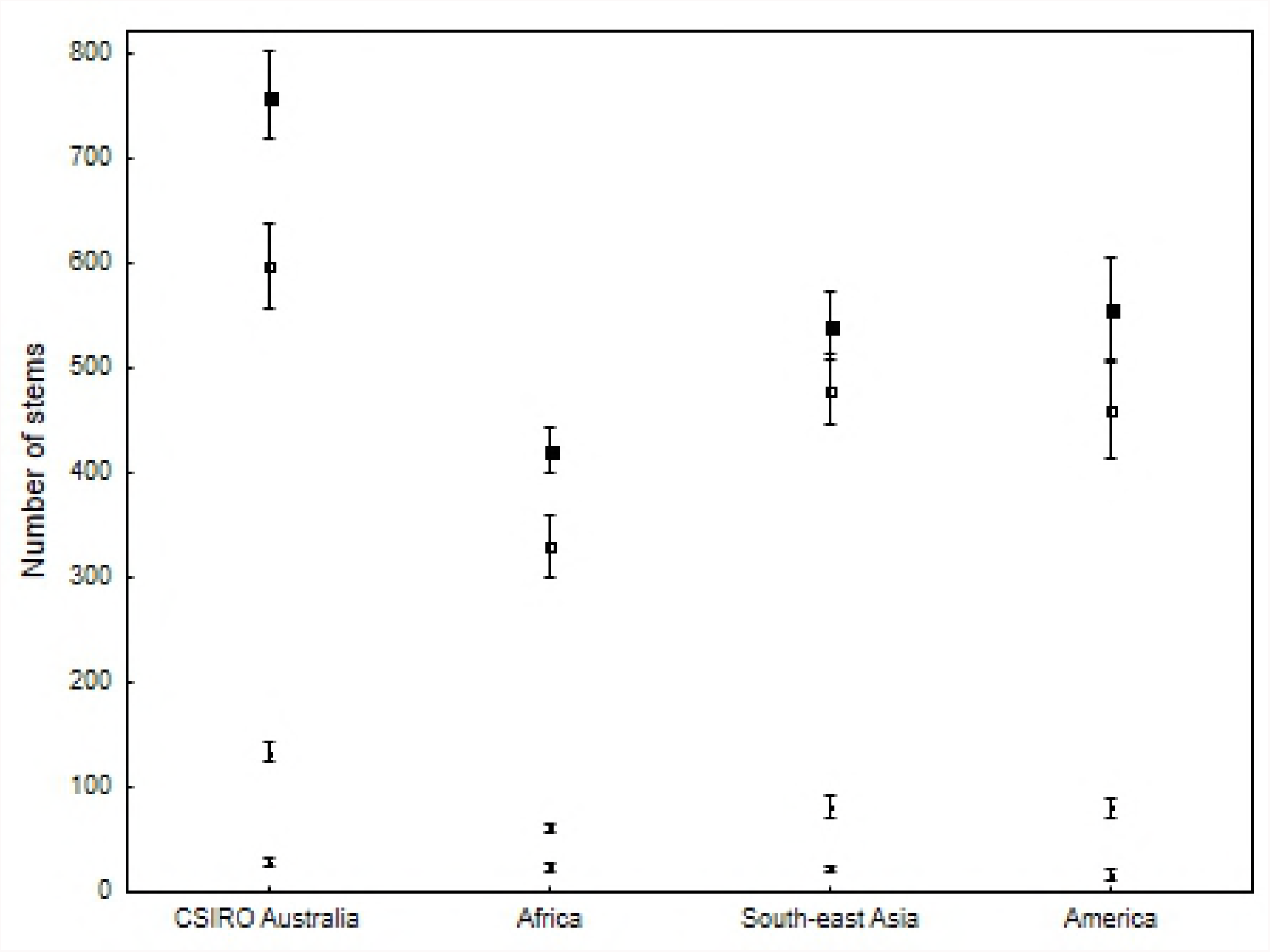
A cross continental comparison of the number of trees per hectare ≥10 cm DBH. From top to bottom; all stems ≥10 cm DBH, stems 10 – 30 cm, stems 30 – 60 cm, stems >60 cm. CSIRO Australia plots have significantly more stems in each comparisons except >60 cm DBH. Data is taken from rainforest plots >10 ha in size, <18.0^0^ north and south of the equator, and within the elevational range of the CSIRO plots. Africa; 6 plots; [24, 43, 44], south-east Asia; 7 plots; [45-50], America; 8 plots: [51-54]. Stems >60 cm DBH are considered large due to the availability of relevant comparative data.

Neither tree height, wood density, nor soil fertility appear to play a significant role in contributing to higher biomass in Australian tropical forests. Mean canopy heights in Australian wet tropical rainforest are in the order of 20-35 m [27, 35], emergent trees are rare, and asymptotic maximum tree heights are similar to America and significantly lower than those in Asia and Africa [55]. The mean wood density for species in our study is 0.63 g/cm^3^, (± 0.16 SD, n = 443) with the most frequent density class being 0.6-0.7 g/cm^3^. This places wood density of Australian species higher than reported means for tropical Africa (0.50 g/cm^3^), Asia (0.57 g/cm^3^), and America (0.60 g/cm^3^) all having the most frequent density class of 0.5-0.6 g/cm^3^ [56]. However, applying slightly higher wood density values to AGB estimations is unlikely to account for the high AGB in Australian wet tropical rainforest particularly as we show that larger species have significantly lower wood densities. There is no relationship between soil fertility and high AGB across our plots. Only one plot is considered eutrophic (measured using exchangeable Ca and Total P%); ep33 of recent volcanic origin with an AGB of 806 Mg/ha. Fifteen of the remaining plots are on highly weathered soils and are considered oligotrophic. Of these, ep18 and ep3 are ranked the second and third highest AGB. This broadly contradicts a global pattern of increasing AGB with increasing soil fertility identified by Slik et al. [4].

## Conclusion

We demonstrate that the contribution to AGB by the largest trees in Australian wet tropical rainforest is generally consistent with that shown for tropical rainforest globally although the high AGB and high contribution from smaller stems introduces some uncertainty in predicting AGB from these large trees. We show that in stark contrast to African and Amazonian forest, our forests have low biomass hyperdominance of larger trees. This puts them in a favourable position to withstand effects of environmental change or large scale disturbance events. Finally, the high average AGB in Australian tropical forests is driven primarily by the relatively high density of large trees coupled with contributions from the higher densities of medium size trees.

## Acknowledgements

We thank Suzanne Prober and Garry Cook (CSIRO) and two anonymous reviewers for helpful suggestions in revising the manuscript. We acknowledge the foresight of Dr Geoff Stocker in establishing the CSIRO permanent plots and the many CSIRO staff and volunteers who have helped to measure and maintain the plots since 1971. The work was carried out under various permits issued by the Queensland Department of Environment and Science.

## Supporting Information

**S1 Fig The relationship between measured tree height from 1998 CSIRO plot data and derived tree height.** a) height derived from [34] equation 6a (grey circles; y = 3.444 + 0.477*x; r^2^ = 0.6285) and, b) height derived from [31] for Australian moist forest (back circles; y = 5.284 + 0.7766*x; r^2^ = 0.6293).

**S2 Fig The relationship between measured tree height from Robson Creek 25 ha plot data [27] and derived tree height.** a) height derived from [34] equation 6a (grey circles; y = 3.2759 + 0.478*x; p = 0.0000; r^2^ = 0.6958 and, b) height derived from [31] for Australian moist forest (back circles; y = 5.4336 + 0.6638*x; p = 0.0000; r^2^ = 0.6866).

**S3 Fig Relationship between estimated AGB using measured height, estimated AGB using derived height and estimated AGB using [33] with no height.** a) estimated AGB using [34] equation 2 with height derived from [34] equation 6a (triangles, dashed fit; y = 20.4771 + 0.6152*x; p = 0.0000; r^2^ = 0.9462), b) estimated AGB using [34] equation 2 using height derived from [31] for Australian moist forests (grey circles and fit; y = −59.9526 + 1.2396*x; r = 0.9793, p = 0.0000; r^2^ = 0.9590). c) estimated AGB using [33] for moist forests without height (black circles and fit; y = 1.2481 + 1.1607*x; r = 0.9852, p = 0.0000; r^2^ = 0.9706).

